# Metabolomics and proteomics analyses of grain yield reduction in rice under abrupt drought-flood alternation

**DOI:** 10.1101/271940

**Authors:** Qiangqiang Xiong, Xiaorong Chen, Tianhua Shen, Lei Zhong, Changlan Zhu, Xiaosong Peng, Xiaopeng He, Junru Fu, Linjuan Ouyang, Jianmin Bian, Lifang Hu, Xiaotang Sun, Jie Xu, Dahu Zhou, Huiying Zhou, Haohua He

**Author notes:** Email: Qiangqiang Xiong –; Xiaorong Chen – Tianhua Shen –; Lei Zhong –; Changlan Zhu –; Xiaosong Peng –; Xiaopeng He –; Junru Fu –; Linjuan Ouyang –; Jianmin Bian –; Lifang Hu –; Xiaotang Sun –; Jie Xu –; Dahu Zhou –; Huiying Zhou –; Haohua He –.

## Abstract

**Highlight:** Abrupt drought-flood alteration is a frequent meteorological disaster that occurs during summer in southern China and the Yangtze river basin, which often causes a large area reduction of rice yield. We previously reported abrupt drought-flood alteration effects on yield and its components, physiological characteristics, matter accumulation and translocation, rice quality of rice. However, the molecular mechanism of rice yield reduction caused by abrupt drought-flood alternation has not been reported.

In this study, four treatments were provided, no drought and no floods (control), drought without floods (duration of drought 10 d), no drought with floods (duration of floods 8 d), and abrupt drought-flood alteration (duration of drought 10 d and floods 8 d). The quantitative analysis of spike metabolites was proceeded by LC-MS (liquid chromatograph-mass spectrometry) firstly. Then the Heat-map, PCA, PLS-DA, OPLS-DA and response ranking test of OPLS-DA model methods were used to analysis the function of differential metabolites (DMs) during the rice panicle differentiation stage under abrupt drought-flood alteration. In addition, relative quantitative analysis of spike total proteins under the treatment was conducted iTRAQ (isobaric tags for relative and absolute quantification) and LC-MS. In this study, 5708 proteins were identified and 4803 proteins were quantified. The identification and analysis of DEPs function suggested that abrupt drought-flood alteration treatment can promote carbohydrate metabolic, stress response, oxidation-reduction, defense response, and energy reserve metabolic process, etc, during panicle differentiation stage. In this study relative quantitative proteomics, metabolomics and physiology data (soluble protein content, superoxide dismutase activity, hydrogen peroxidase activity, peroxidase activity, malondialdehyde content, free proline content, soluble sugar content and net photosynthetic rate) analysis were applied to explicit the response mechanism of rice panicle differentiation stage under abrupt drought-flood alteration and provides a theoretical basis for the disaster prevention and mitigation.

**Abstract:** Abrupt drought-flood alternation is a meteorological disaster that frequently occurs during summer in southern China and the Yangtze river basin, often causing a significant loss of rice production. In this study, a quantitative analysis of spike metabolites was conducted via liquid chromatograph-mass spectrometry (LC-MS), and Heat-map, PCA, PLS-DA, OPLS-DA, and a response ranking test of OPLS-DA model methods were used to analyze functions of differential metabolites (DMs) during the rice panicle differentiation stage under abrupt drought-flood alternation. The results showed that 102 DMs were identified from the rice spike between T1 (abrupt drought-flood alternation) and CK0 (control) treatment, 104 DMs were identified between T1 and CK1 (drought) treatment and 116 DMs were identified between T1 and CK2 (flood) treatment. In addition, a relative quantitative analysis of spike total proteins was conducted using isobaric tags for relative and absolute quantification (iTRAQ) and LC-MS. The identification and analysis of DEPs functions indicates that abrupt drought-flood alternation treatment can promote carbohydrate metabolic, stress response, oxidation-reduction, defense response, and energy reserve metabolic process during the panicle differentiation stage. In this study, relative quantitative metabolomics and proteomics analyses were applied to explore the response mechanism of rice panicle differentiation in response to abrupt drought-flood alternation.

**Abbreviations:**

CK0: no drought and no floods
CK1: drought without floods
CK2: no drought with floods
T1: abrupt drought-flood alteration
LC-MS: liquid chromatograph-mass spectrometry
PCA: principle component analysis
(O)PLS-DA: (orthogonal) partial least-squares-discriminant analysis
DMs: differential metabolites
iTRAQ: isobaric tags for relative and absolute quantification
DEPs: differentially expressed proteins
KEGG: kyoto encyclopedia of genes and genomes
GO: gene ontology
SOD: superoxide dismutase
CAT: hydrogen peroxidase
POD: peroxidase
MDA: malondialdehyde
Pn: net photosynthetic rate
ROS: reactive oxygen species
VIP: variable importance in the projection
FC: fold change

## Introduction

Rice (*Oryza sativa*) is one of the prominent food crops globally and is considered a major staple food in many Asian countries; here, each person consumes more than 100 kg of rice per year, on average (Nelson, 2011). Since the green revolution in 1960, there has been a progressive increase in rice yield. However, the ever-growing population and adverse climatic developments pose enormous challenges for a sustainable meeting of the rice demand, as its growth and yield are greatly influenced by drought and flooding stress (Sarvestani *et al.*, 2008; Gautam *et al.*, 2015). This issue will remain a severe threat to food security unless means to circumvent the impacts of water stress can be developed. Rice plants have developed the ability to adapt to environmental changes during the long-term acclimation process, such as changes in water availability. At the same time, floods during spring, during the succession of spring to summer in the Yangtze river basin, and floods during the succession of summer to autumn occur frequently in southern China, e.g. in Jiangxi, Hunan, Anhui, and Guangdong provinces of China. This usually greatly reduces rice yield. Meteorological data shows that drought and flood have increased in severity over the past 30 years in Jiangxi and other southern regions (Cai *et al.*, 2013). In particular, drought conditions followed by flood over a large area significantly reduces the rice yield, which has been called “abrupt drought-flood alternation”. This abrupt drought-flood alternation is characterized by a persistent drought during the early stage and is then followed by flooding weather (Wang *et al.*, 2009). Particularly during recent years, the abrupt drought-flood alternation occurrence trend became increasingly frequent, causing a reduction of crop yield in large areas, even resulting in the complete loss of crop yield; therefore, it has attracted much attention (Wang *et al.*, 2009; Peng *et al.*, 2009; Deng and Chen, 2013). According to statistical data obtained in previous studies, abrupt drought-flood alternation occurred over 14 years between 1960 and 2012 in China. Once every four years, and a total of 23 extreme climate conditions of abrupt drought-flood alternation occurred at the Huai river (Wang *et al.*, 2014). In 2011, 42 counties (cities or districts), encompassing nearly three million people and a crop area of 9.04 million hm^2^, were affected by such an event, which led to an economic loss of 5.8 billion Yuan for Jiangxi province. Especially in the Huaibei plain area, many severe abrupt drought-flood alternation natural disasters have occurred (Shen *et al.*, 2012). Furthermore, it was found that compared to a control treatment, the yield of abrupt drought-flood alternation treatment was reduced by 30.3%. This was most unfavorable for the yield, which became apparent through an analysis on the yield reducion law of rice under the conditions of abrupt drought-flood alternation (Gao *et al.*, 2017).

To date, some studies have focused on problems associated with cultivation, physiology, ecology, genetics, breeding, and other aspects of damage of rice caused by drought (Kumar *et al.*, 2014; Lauteri *et al.*, 2014; Majeed *et al.*, 2013; Hadiarto and Tran, 2011) and flooding (Dwivedi *et al.*, 2017; Winkel *et al.*, 2014; Singh *et al.*, 2014; Gautam *et al.*, 2017). Abrupt drought-flood alternation affects yield and its components, physiological characteristics, matter accumulation and translocation, and rice quality (Zhong *et al.*, 2016; Deng *et al.*, 2017; Xiong et *al.*, 2017a; Xiong *et al.*, 2017b; Xiong *et al.*, 2017c). However, the molecular mechanism with which rice yield reduction is caused by abrupt drought-flood alternation has rarely been reported.

Metabolomics and proteomics became powerful tools for analyzing plant reactions to various environmental stimuli (Benevenuto *et al.*, 2017). Especially comparative studies of the molecular mechanism that underlie plant responses to subjection to adverse conditions (such as drought and flooding) provide valuable insights into plant responses to pre-determined stresses and provides information on the biochemical pathways that participate in the acclimation to environmental constrains (Tuberosa and Salvi, 2006). At present, most studies focus on proteomics and metabolomics at the rice seedling stage to explain effects of drought or flood on rice (Fukao, 2006; Shu *et al.*, 2011; He *et al.*, 2011; Jayaweera *et al.*, 2016; Alpuerto, 2016). In such studies, abundantly identified proteins and metabolites are usually involved in defense mechanisms, including detoxification enzymes, redox status regulation, signaling pathways, protein folding and degradation, photosynthesis, and primary metabolism. However, studies that utilize a combination of proteomics and metabolomics, thus clarifying drought and flooding responses, especially the reasons for yield loss in rice during the panicle differentiation stage under abrupt drought-flood alternation, have not been reported. On the other hand, a large number of super hybrid rice varieties have been cultivated in the Yangtze river basin double-cropping rice of China during recent years. The productive potential level of super rice varieties of has been greatly improved recently, while also increasing the demand for water and fertilization (Zeng *et al.*, 2009; Chang *et al.*, 2014). Therefore, it is necessary to combine proteomics, metabolomics, and physiology to clarify the abrupt drought-flood alternation response mechanism of rice yield reduction during the panicle differentiation stage.

## Materials and methods

### Plant materials and growth conditions

Wufengyou 286 (*Oryza sativa* L.) is the dominant double-cropping super hybrid early rice variety, which was planted in the Jiangxi province, located in a region where double-cropping early rice is conducted in southern China (Wufeng A/Zhonghui 286, a super rice variety certified by the Chinese ministry of agriculture in 2015). The experiment was conducted at the science and technology park of Jiangxi agricultural university (28°46′N, 115°50′E, altitude: 48.8 m, annual average temperature: 17.5°C, average annual sunshine: 1720.8 h, annual average evaporation: 1139 mm, and average annual rainfall: 1747 mm), Nanchang, Jiangxi province of China in 2017. Rice was planted in plastic buckets of 24.0 cm height and 29.0 cm inner diameter of the upper portion, and 23.5 cm inner diameter at the bottom. Soil samples were extracted from the upper soil layer (0 – 20 cm) of the rice experiment field at the experimental site. The physical and chemical properties of the experimental soil are shown in Table 1. The soil was naturally air-dried, pulverized using a soil disintegrator (FT-1000A, Changzhou WIK Instrument Manufacturing Co. Ltd., China), and sieved through a 100-mm mesh. Each pot contained approximately 10 kg of dry soil, which was soaked in water two weeks prior to transplantation, and 5 g compound fertilizer (N-P-K = 15%-15%-15%) was applied to each pot as base fertilization. After one week, 2 g potassium chloride and 2 g urea fertilizer were applied. Seed dormancy was broken via exposure to the sun for 1 d, followed by pre-germination and then, sown in a rice field on 16 March, 2017. Forty-day-old seedlings were then transplanted on 26 April into plastic buckets with only one seedling per hill (three plants per plastic bucket). Insecticides were used to prevent insect damage. All other agronomic practices referred to the local recommendations to avoid yield loss.

**Table 1.** Physical and chemical properties of the experimental soil

| Soil pH | Organic matter<br>(mg kg<sup>-1</sup>) | Total nitrogen<br>(g kg<sup>-1</sup>) | Available nutrient (mg kg<sup>-1</sup>) |  |  |
| --- | --- | --- | --- | --- | --- |
|  |  |  | N (nitrogen) | P (phosphorus) | K (potassium) |
| 5.44 | 26.1 | 1.24 | 110 | 20.8 | 80.5 |

### Treatments

Via analysis of historical data of abrupt drought-flood alteration events (Cheng *et al.*, 2012), it was found that the abrupt drought-flood alteration often occurred in mid and late June in the Yangtze river basin. At this time, the double-cropping early rice of the Yangtze river basin was at the panicle differentiation stage. Therefore, the rice abrupt drought-flood alteration stage was set at the panicle differentiation stage in this experiment. The start and end dates of both drought and flood were set to mimic natural drought and flooding disasters that happened to double-cropping early rice in the Yangtze river basin. For drought treatment, the remaining water in plastic buckets was drained and buckets were moved to a rain shelter prior to rain. The soil moisture content was monitored via vacuum meter type soil moisture meter (measuring range 0 ~ 85 kPa, institute of Chinese academy of sciences), and rice was naturally dried. The soil water potential after drought treatment for 8 d exceeded the maximum measurable value of the instrument, and drought treatment continued for further 2 d until the soil was white and cracking, and the plants were wilting and withered (imitating severe drought). For submergence treatment, plants in soil-containing pots were completely submerged in a high water-filled square box (1.35 m height) in a greenhouse. The water in the high square box was static, clean tap water and was not exchanged during the entire submergence period. For abrupt drought-flood alteration treatment, immediately after the onset of the change from drought treatment to flood treatment, plants of the control treatment (CK0) were maintained in a 3~5 cm water layer. Four treatments (each treatment with three replicates) were applied, and 15 pots were used per replication, using a randomized block design (Table 2).

**Table 2.** Different water treatments for super hybrid early rice (Wufengyou 286)

| Treatments | Duration of drought (d) | Start and end dates (m-d) | Duration of flood (d) | Start and end dates (m-d) |
| --- | --- | --- | --- | --- |
| CK0 | 0 |  | 0 |  |
| CK1 | 10 | June 12–June 22 | 0 |  |
| CK2 | 0 |  | 8 | June 14–June 22 |
| T1 | 10 | June 4–June 14 | 8 | June 14–June 22 |

CK0: no drought and no flood; CK1: drought without flood; CK2: no drought without flood; T1: abrupt drought-flood alteration.

### Yield and physiological parameters

Once rice was mature, 10 undamaged plants per treatment were used to obtain the grain yield per plant. The reciprocal second leaf sample of rice was taken as 0.1 g at the first day after treatment. All samples were flash-frozen in liquid nitrogen and stored at −80°C until measurement. Soluble protein content, superoxide dismutase (SOD) activity, hydrogen peroxidase (CAT) activity, peroxidase (POD) activity, malondialdehyde (MDA) content, free proline content, and soluble sugar content were determined with the kit of suzhou kemin biotechnology Co., Ltd., China (Micro method).

Net photosynthetic rate (Pn) was measured at the reciprocal second leaves of rice on June 22 (the first day after treatment), from 9:30 to 12:00 am. Three biological replicates were performed per treatment. Pn was measured using a CI-340 portable photosynthesis analyzer (CID Bio-Science, USA).

### Metabolite extraction and analysis

The spikes of three plants were sampled after drought treatment, flood treatment, and abrupt drought-flood alteration treatment, respectively. Eight biological replicates were performed per treatment sample (each replicate contained three plants). All samples were flash-frozen in liquid nitrogen and stored at −80°C until metabolite extraction. Samples were ground in liquid nitrogen with a pestle and mortar. Separate aliquots of the frozen and ground sample were used for metabolite extraction as previously described (Bowne *et al.*, 2012; Zhao *et al.*, 2013). The extracted samples were then subjected to derivatization and analyzed via liquid chromatograph-mass spectrometry (LC-MS). The LC-MS system comprised AN ACQUITY UHPLC system (Waters Corporation, Milford, USA) coupled with an AB SCIEX Triple TOF 5600 System (AB SCIEX, Framingham, MA), which was used to analyze the metabolic profiling in both ESI positive and ESI negative ion modes. An ACQUITY UPLC BEH C18 column (1.7 μm, 2.1 × 100 mm) was employed in both positive and negative modes. Data acquisition was performed in full scan mode (m/z ranges from 70 to 1000) combined with IDA mode. QC samples were prepared by mixing aliquots of all samples into a pooled sample. The QCs were injected at regular intervals (every 10 samples) throughout the analytical run to provide a data set from which repeatability can be assessed.

### Metabolite data preprocessing and statistical analysis

The raw data were converted to common data format files (mzML) using the conversion software program MSconventer. Metabolomics data were acquired using the software XCMS, version 1.50.1, which produced a matrix of features with their associated retention time, accurate mass, and chromatographic data. Variables that presented at least 50% of either group were extracted. Isotope and internal standard were removed from the data set. To achieve minimum RSD, all ions were normalized to the total peak area of each sample in Excel 2007 (Microsoft, USA). 1986 metabolite ions were acquired in positive ion mode, while 899 metabolite ions were acquired in negative ion mode. 92.48% of the ions in positive ion mode and 89.52% of those in negative ion mode exhibited less than 30% of RSD, thus displaying good reproducibility of the metabolomics method. The metabolite ions with RSD% below 30% were used for further data processing.

Positive and negative data were combined to form a combined data set, which was then imported into the SIMCA software package (version 14.0, Umetrics, Umeå, Sweden). Principle component analysis (PCA) and (orthogonal) partial least-squares-discriminant analysis (O)PLS-DA were conducted to visualize metabolic alterations among experimental groups, following mean centering and unit variance scaling. The Hotelling’s T2 region, shown as an ellipse in score plots of the models, defined the 95% confidence interval of the modeled variation. Variable importance in the projection (VIP) ranked the overall contribution of each variable to the OPLS-DA model, and those variables with VIP > 1 were considered relevant for group discrimination. In this study, the default 7-round cross-validation was applied where 1/seventh of the samples were excluded from the mathematical model per round, to avoid overfitting.

### Identification of differential metabolites (DMs)

DMs were chosen on the basis of the combination of a statistically significant threshold of variable influence on projection (VIP) values obtained from the OPLS-DA model. Also, *p*-values from a two-tailed Student’s t test on the normalized peak areas were used, where metabolites with VIP values above 2, with fold change (FC) above 1.75, and *p*-values below 0.005 were included, respectively. We used the software One-step solution to identify small molecules in metabolomics studies, which was co-developed with the Dalian institute of chemical physics, at the Chinese academy of sciences and Dalian chem data solution information technology Co., Ltd to identify DMs. HMDB and METLIN reference material databases (built by the Dalian Institute of chemical physics) were used.

### Protein extraction and peptide preparation

The spikes of three plants were sampled after drought treatment, flood treatment, and abrupt drought-flood alteration treatment, respectively; two replicates were performed per treatment sample (each replicate had three plants). All samples were flash-frozen in liquid nitrogen and stored at −80°C for protein extraction. The protein extract was digested according to the filter-aided sample preparation procedure (Wiśniewski *et al.*, 2009). After trypsin digestion, peptides were desalted via Strata X C18 SPE column (Phenomenex) and vacuum-dried. Peptides were reconstituted in 0.5 M TEAB and peptide mixtures were labeled via iTRAQ-8plex, processed according to the manufacturer’s protocol for the iTRAQ kit. Briefly, one unit of iTRAQ reagent (AB Sciex, Foster City, CA, USA) was thawed and reconstituted in acetonitrile. The peptide mixtures were then incubated for 2 h at room temperature and pooled, desalted, and dried via vacuum centrifugation. The tryptic peptides were fractionated into fractions via high pH reverse-phase HPLC using Agilent 300Extend C18 column (5 μm particles, 4.6 mm ID, and 250 mm length). Briefly, peptides were first separated with a gradient from 8% to 32% acetonitrile (PH 9.0) over 60 min into 60 fractions. Then, the peptides were combined into 18 fractions and dried via vacuum centrifugation.

### Peptides LC-MS/MS analysis and database search

The tryptic peptides were dissolved in 0.1% formic acid (solvent A) and directly loaded onto a home-made reversed-phase analytical column (15 cm length and 75 μm i.d.). The gradient comprised an increase from 6% to 23% solvent B (0.1% formic acid in 98% acetonitrile) over 26 min, 23% to 35% during 8 min, then increasing to 80% during 3 min, and then holding at 80% for the last 3 min, all at a constant flow rate of 400 nL/min on an EASY-nLC 1000 UPLC system.

The peptides were subjected to NSI source followed by tandem mass spectrometry (MS/MS) in Q Exactive™ Plus (Thermo), which was coupled online to the UPLC. The applied electrospray voltage was 2.0 kV. The m/z scan range was 350 to 1800 for full scan, and intacting peptides were detected in the Orbitrap at a resolution of 70,000. Peptides were then selected for MS/MS using the NCE setting at 28 and the fragments were detected in the Orbitrap at a resolution of 17,500. A data-dependent procedure that alternated between one MS scan followed by 20 MS/MS scans with 15.0 s dynamic exclusion. Automatic gain control (AGC) was set to 5E4. The fixed first mass was set to 100 m/z.

The resulting MS/MS data were processed using the Maxquant search engine (v.1.5.2.8). Tandem mass spectra were searched against the 39946_Oryza sativa indica database and were concatenated with reverse decoy database. Trypsin/P was specified as cleavage enzyme, allowing up to two missing cleavages. The mass tolerance for precursor ions was set to 20 ppm for the First search and to 5 ppm for the Main search, and the mass tolerance for fragment ions was set to 0.02 Da. Carbamidomethyl on Cys was specified as fixed modification and oxidation on Met was specified as variable modification. FDR was adjusted to < 1% and the minimum score for peptides was set to > 40.

### Bioinformatics analysis

The Gene Ontology (GO) annotation proteome was derived from the UniProt-GOA database (www. http://www.ebi.ac.uk/GOA/). Firstly, identified protein ID was converted to an UniProt ID and then mapped to GO IDs via the protein ID. If some identified proteins were not annotated by the UniProt-GOA database, the InterProScan soft would be used to annotate the protein’s GO functional based on the protein sequence alignment method. Then, proteins were classified by Gene Ontology annotation, based on three categories: biological process, cellular component, and molecular function. For each category, a two-tailed Fisher’s exact test was employed to test the enrichment of the differentially expressed protein against all identified proteins. GO with a corrected *p*-value < 0.05 was considered significant.

The Kyoto Encyclopedia of Genes and Genomes (KEGG) connects known information of molecular interaction networks, such as pathways and complexes (the “Pathway” database), information about genes and proteins generated by genome projects (including the gene database), and information about biochemical compounds and reactions (including compound and reaction databases). Such databases are different networks, known as the “protein network”, and the “chemical universe”, respectively. Efforts are in progress to add to the knowledge of KEGG, including information regarding ortholog clusters in the KEGG Orthology database. KEGG Pathways mainly include: metabolism, genetic information processing, environmental information processing, cellular processes, rat diseases, and drug development. The KEGG database was used to annotate the protein pathway. Firstly, the KEGG online service tool KAAS (KEGG automatic annotation server) was used to annotate the protein’s KEGG database description. Then, the annotation result was mapped on the KEGG pathway database using the KEGG online service tool KEGG mapper. The KEGG database was used to identify enriched pathways via two-tailed Fisher’s exact test to test the enrichment of the differentially expressed proteins against all identified proteins. The pathway with a corrected *p*-value < 0.05 was considered significant. These obtained pathways were classified into hierarchical categories according to the KEGG website.

For further hierarchical clustering based on different protein functional classification (such as: GO, pathway, or complex), we first collated all obtained categories after enrichment along with their *p*-values, and then filtered for those categories that were at least enriched in one of the clusters with a *p*-value < 0.05. This filtered *p*-value matrix was transformed with the function x = −log_10_ (*p*-value). Finally, these x values were z-transformed for each functional category. These z scores were then clustered via one-way hierarchical clustering (Euclidean distance and average linkage clustering) in Genesis. Cluster memberships were visualized via a heat map using the “heatmap.2” function of the “gplots” R-package.

### Statistical analysis

Origin 8.5 was used to arrange and map the yield as well as both physiological and biochemical data. SPSS 17.0 software was used to conduct analyses of variance (ANOVA) and Tukey tests for multiple comparisons and significant analysis.

## Results

### Grain yield and physiological indexes

After the drought, flood, and abrupt drought-flood alternation during panicle differentiation stage, and compared to CK0, grain yield per plant decreased significantly (*P* < 0.01). No significant difference was found between CK1, CK2, and T1 (*P* > 0.05); however, grain yield per plant was lowest at T1. Compared to CK1 and CK2, grain yield per plant of T1 decreased by 17.22% and 34.90%, respectively (Fig. 1a). Compared to CK0 and CK1, the soluble protein content of T1 decreased significantly (*P* < 0.01), and no significant difference was found with CK2 (Fig. 1b). The SOD activity of T1 was significantly higher compared to CK0 and CK1, and the difference was significant (*P* < 0.01); however, there was no significant difference between T1 and CK2 (*P* > 0.05; Fig. 1c). The CAT activity of T1 was significantly higher compared to CK0 and CK2 (*P* < 0.01); however, it was significantly lower compared to CK1 (*P* < 0.01; Fig. 1d). The POD activity of T1 was highest, and significantly differed from those of CK0, CK1, and CK2 (*P* < 0.01), indicating that the effect of abrupt drought-flood alternation on POD was significant (Fig. 1e). The MDA content of CK1 was highest, and there was a significant difference between CK1 and T1 (*P* < 0.01). The MDA content of T1 was 10.37% higher than that of CK0; however, there was no significant difference (*P* > 0.05; Fig. 1f). Compared to CK0 and CK1, the free proline content and soluble sugar content of T1 decreased significantly (*P* < 0.01) and there was no significant difference to CK2 (*P* > 0.05) (Fig. 1g, h). The net photosynthetic rate of T1 was the lowest of all treatments. Compared to CK0, the net photosynthetic rate of T1 decreased significantly (*P* < 0.01). Compared to CK1 and CK2, the net photosynthetic rate of T1 decreased by 3.42% and 12.87%, respectively. This indicated that drought and flooding, especially abrupt drought-flood alternation was strongly detrimental to the photosynthetic capacity of a rice leaf.

**Figure 1.**
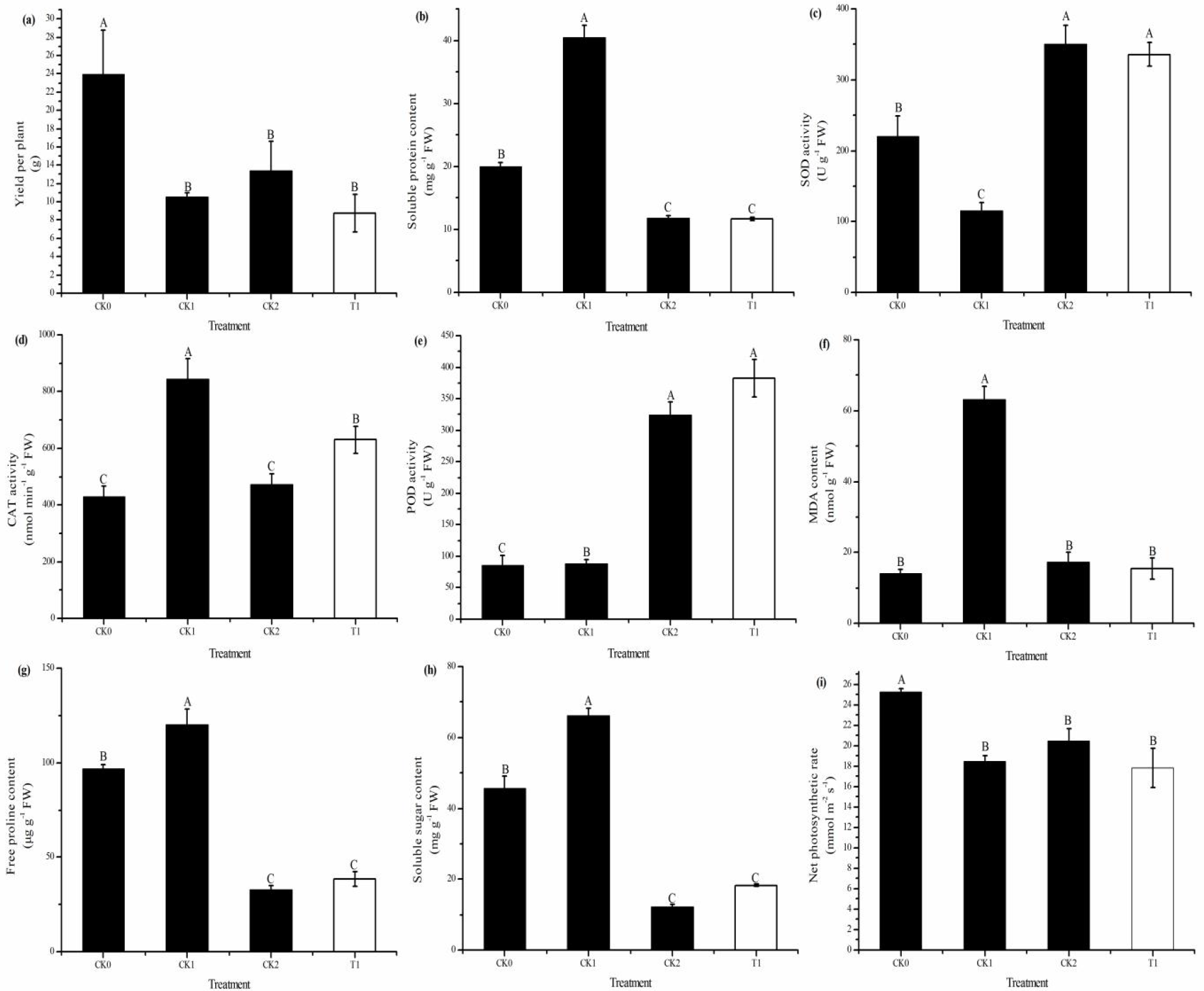
Analysis of yield and physiological indexes: (a) yield per plant, (b) soluble protein content, (c) SOD activity, (d) CAT activity, (e) POD activity, (f) MDA content, (g) free proline content, (h) soluble sugar content, and (i) net photosynthetic rate. The column data in figure shows the average and the short lines represents the mean square deviations. Different capital letters indicate the significance level of differences among treatments at the 0.01 level (*P* < 0.01). CK0: no drought and no floods; CK1: drought without floods; CK2: no drought with floods; T1: abrupt drought-flood alteration.

### Metabolite profiles

Metabolic profiling via LC-MS was performed in an attempt to identify changes in metabolites that result from the differential regulation of the enzymes involved in the above-mentioned processes. In total, 102 metabolites were identified from the rice spike, of which, expressions significantly differed between the T1 and the CK0 treatment; 104 metabolites were identified and were found to be significantly different between the T1 and the CK1 treatment; 116 metabolites were identified and were found to be significantly different between the T1 and the CK2 treatment. Detailed DMs identification information can be found in supplementary Table S1.

Hierarchical cluster analysis showed that the three compared groups segregated separately based on their metabolomic responses. The metabolite profiles of different treatments in response to the abrupt drought-flood alternation treatment were also analyzed via Heatmap (Fig. 2a, b, c). This visualized that abrupt drought-flood alternation treatment responded differently to water induced stresses. Principal component analysis (PCA), partial least squares discriminant analysis (PLS-DA), and orthogonal PLS-DA also segregated T1 from CK0 (Fig. 3a, d, g), CK1 (Fig. 3b, e, h), and CK2 (Fig. 3c, f, i) based on metabolite levels in each compared group.

**Figure 2.**
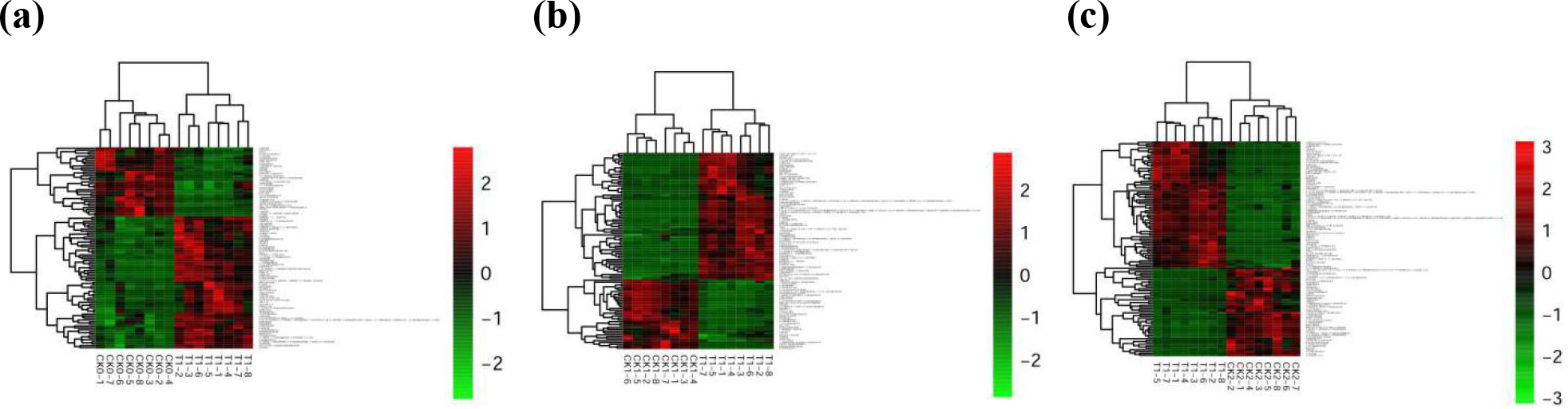
Hierarchical cluster analysis of changed metabolite pools. Hierarchical trees were drawn based on detected changes of metabolites in spikes of rice under different water treatments: (a) T1 vs CK0 comparison treatment, (b) T1 vs CK1 comparison treatment, and (c) T1 vs CK2 comparison treatment. Columns represent the repetition between different treatments, while rows represent different metabolites. Red and green colors indicate increased and decreased metabolite concentrations, respectively. CK0: no drought and no floods; CK1: drought without floods; CK2: no drought with floods; T1: abrupt drought-flood alteration.

**Figure 3.**
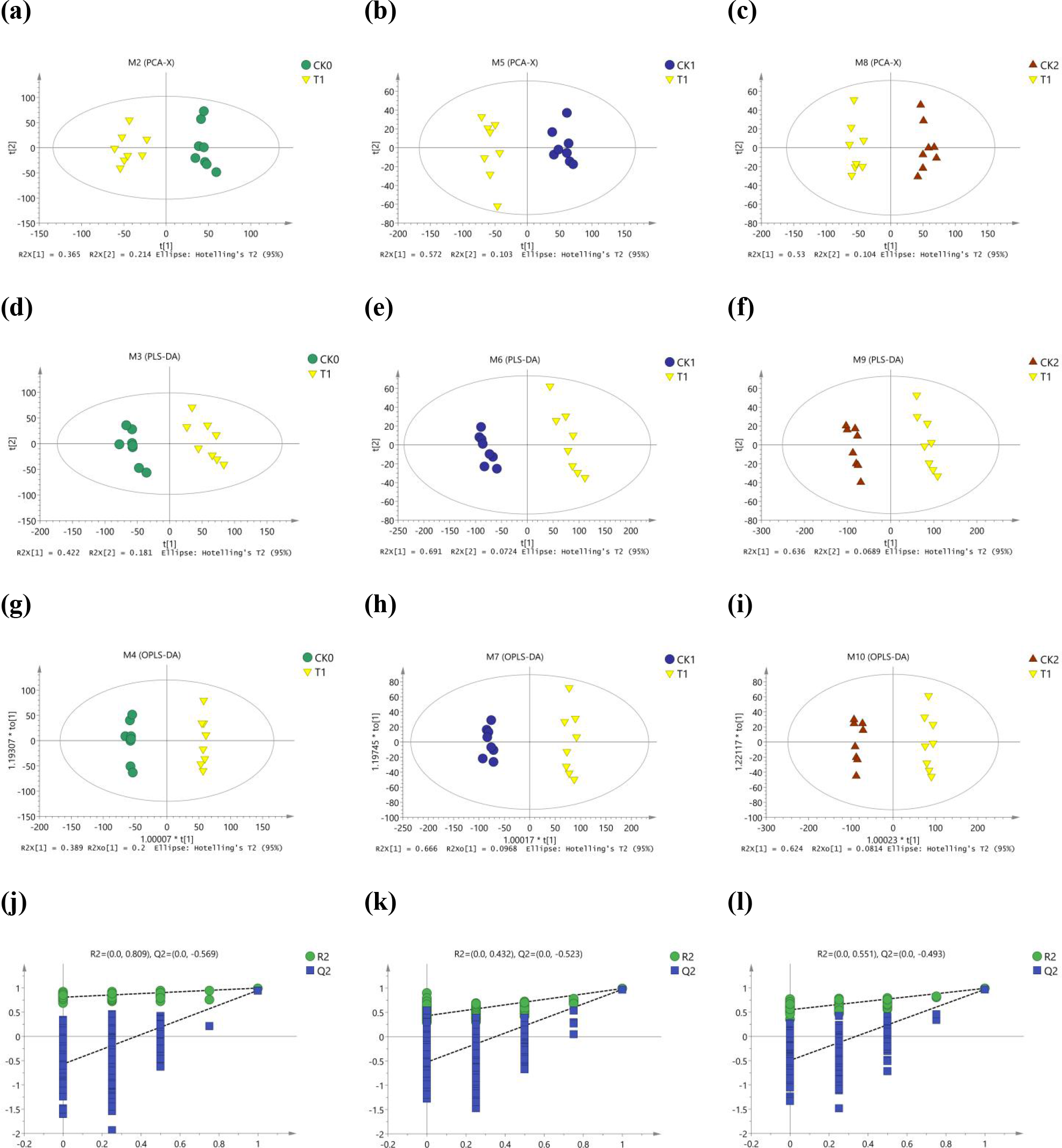
Separation of the different comparison treatments, (a), (b), and (c): PCA; (d), (e), and (f): PLS-DA; (g), (h), and (i): OPLS-DA; (j) T1 vs CK0 comparison treatment, (k) T1 vs CK1 comparison treatment, (l) T1 vs CK2 comparison treatment response sorting test of the OPLS-DA model. CK0: no drought and no floods; CK1: drought without floods; CK2: no drought with floods; T1: abrupt drought-flood alteration.

To test whether model reproducibility is good and whether the data in the model are fitting, it is suggested to avoid the classification obtained by the supervised learning method. We conducted 200 response sorting tests of the OPLS-DA model: fixed X matrix and previously defined Y matrix variable classification (e.g. 0 or 1) were randomly arranged n times (n = 200) and the OPLS-DA model was established to obtain the corresponding stochastic model of *R*^*2*^ and *Q*^*2*^ value. With the original model of *R*^*2*^*Y* and *Q*^*2*^*Y* linear regression, the regression line and the *Y* axis intercept values were *R*^*2*^ and *Q*^*2*^, and were used to measure whether the model was over fitting. When the intercept of *Q*^*2*^ is below zero, the model is valid (Mahadevan et al. 2008). Figures 3j, 3k, and 3l show that the model was still be valid after cross validation.

### Identification of differentially expressed proteins (DEPs)

iTRAQ labeling coupled with LC-MS was utilized to profile the DEPs between CK0, CK1, CK2, and T1. To evaluate the reliability of the data that was generated via proteomic analysis, the Pearson’s correlation coefficient value was calculated, resulting in 0.86, 0.88, 0.86, and 0.87 for CK0-1 and CK0-2, CK1-1 and CK1-1, CK2-1 and CK2-2, and T1-1 and T1-2, respectively. Each treatment was conducted in two replicates; then, heatmaps were drawn via the R-package 3.4.1 (Fig. 4). The obtained heatmaps indicated good reproducibility of two biological replicates in different treatments.

**Figure 4.**
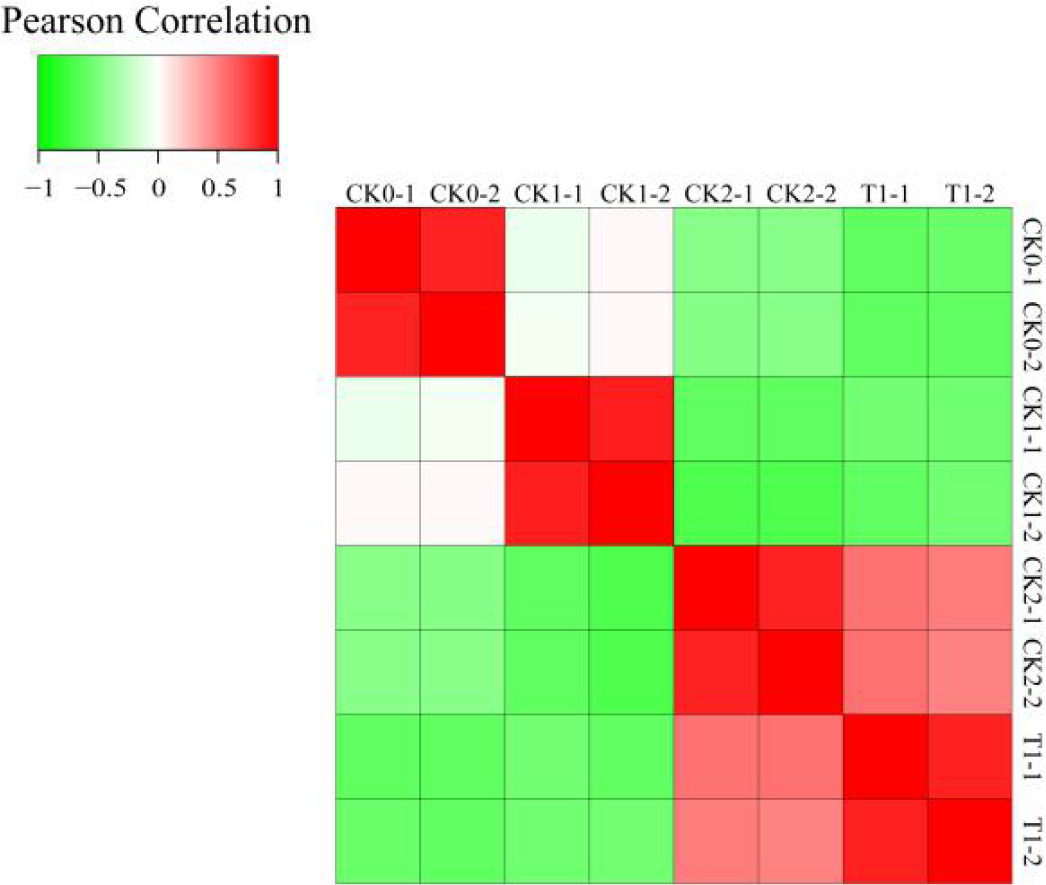
Repeatability test between each two samples. A Pearson coefficient closer to −1 indicates a negative correlation, a coefficient closer to 1 indicates a positive correlation, and a coefficient closer to 0 indicates no correlation. Red, green, and white indicate different degrees of correlation. CK0-1 and CK0-2: no drought and no floods; CK1-1 and CK1-1: drought without floods; CK2-1 and CK2-2: no drought with floods; T1-1 and T1-2: abrupt drought-flood alteration.

In the present study, a total of 199822 spectrums were detected, 48152 of which could be matched to peptides in the database and 23148 were unique peptides. In total, 5708 proteins could be identified and 4803 proteins were experimentally quantified, which included the different treatments and replicates of the iTRAQ proteomic analysis (Table 3). Detailed information about the identification and quantification of these two independent biological replicates in different treatments can be found in the supplementary Excel sheets (sheet1 shows T1 vs CK0, sheet2 shows T1 vs CK1, and sheet3 shows T1 vs CK2).

**Table 3.** MS/MS spetrum database search analysis summary

| Total<br>spectrums | Matched<br>spectrums | Peptides | Unique<br>peptides | Identified<br>proteins | Quantifiable<br>proteins |
| --- | --- | --- | --- | --- | --- |
| 199822 | 48152 | 25602 | 23148 | 5708 | 4803 |

Proteins with a fold-change (FC) > 1.5 or FC < 0.67 and *p* < 0.05 between the treatment (T1) and control groups (CK0, CK1, and CK2) were classified as DEPs, and DEPs were hence considered as abrupt drought-flood alternation at panicle differentiation stage responsive proteins. These cut-offs were selected based on a previous publication that investigated the reproducibility of iTRAQ™ quantification (Moulder *et al.*, 2010). According to this criterion, 522 proteins were found to exhibit significant differential expression between T1 and CK0; 528 proteins were found to exhibit significant differential expression between T1 and CK1; 82 proteins were found to exhibit significant differential expression between T1 and CK2; and the up-regulated and down-regulated DEPs in the different comparison groups are shown in the supplementary Excel sheets (sheet1 shows T1 vs CK0, sheet2 shows T1 vs CK1, and sheet3 shows T1 vs CK2). The three sample comparable group had 54 DEPs in common (Fig. 5).

**Figure 5.**
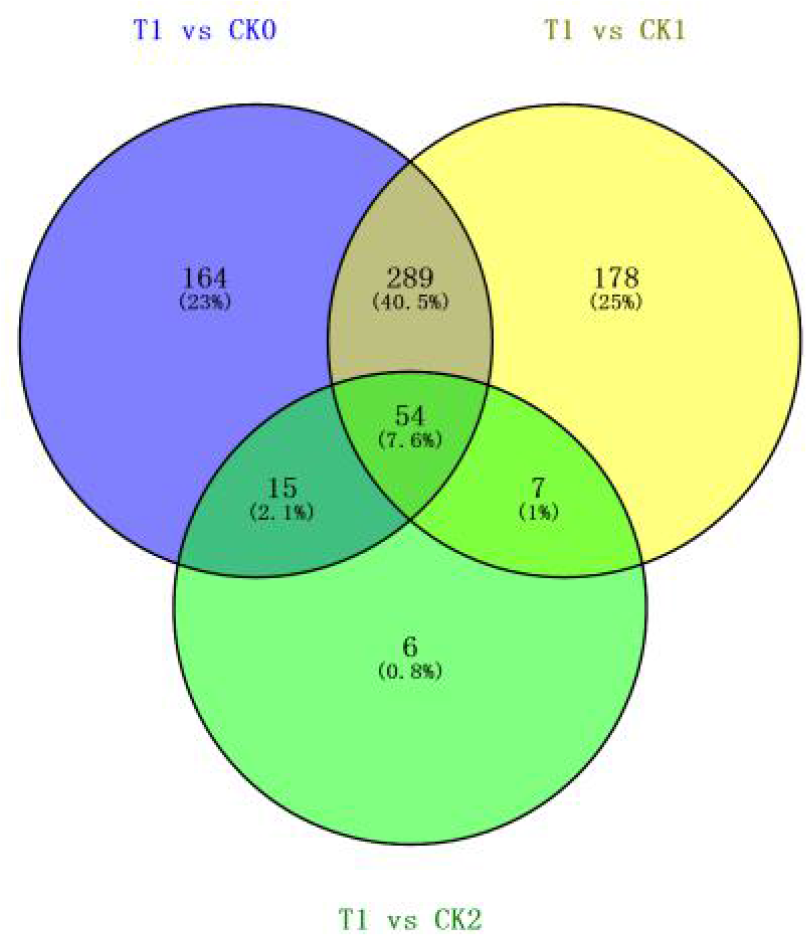
Venn diagram the differentially expressed proteins (DEPs) between the treatment group (T1) and control groups (CK0, CK1, and CK2). Results for abrupt drought-flood alternation at the young spike differentiation stage responsive proteins. CK0: no drought and no floods; CK1: drought without floods; CK2: no drought with floods; T1: abrupt drought-flood alteration.

**Figure 6.**
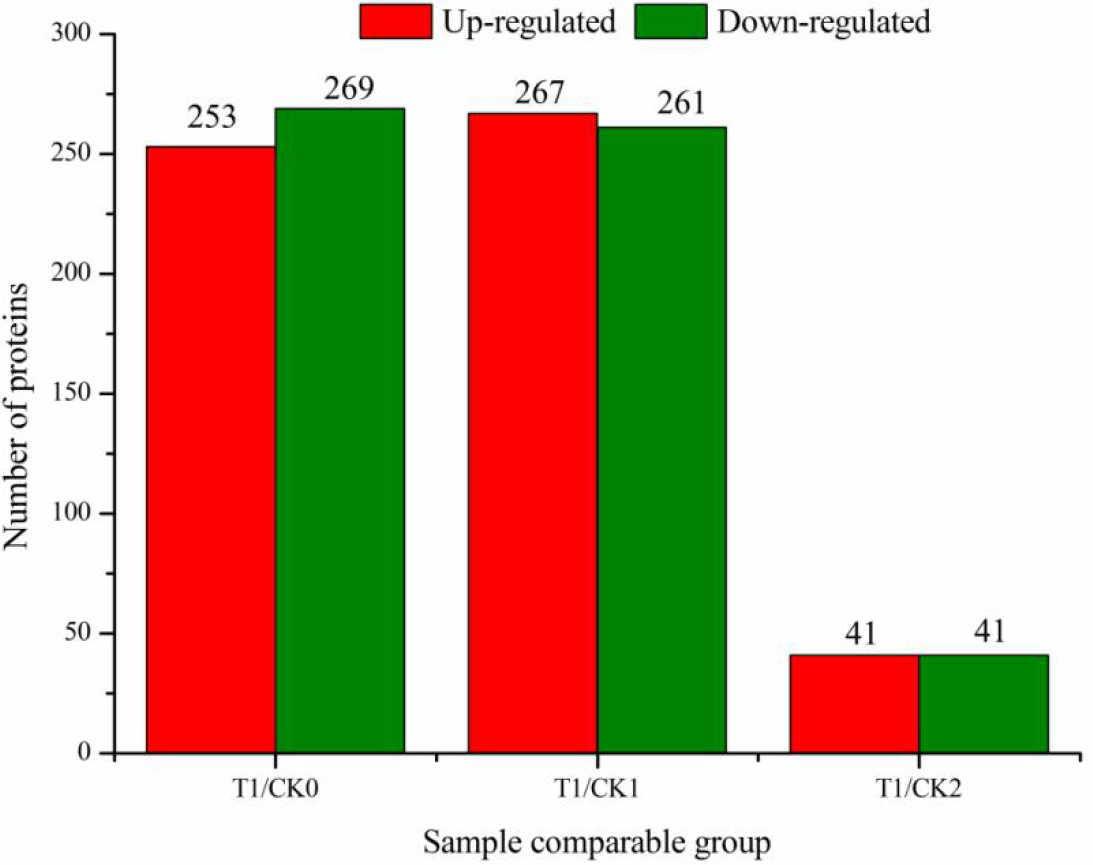
Summary of up- and down-regulation of differentially expressed proteins (DEPs) between the treatment group (T1) and control groups (CK0, CK1, and CK2), DEPs with a fold-change (FC) > 1.5 or FC < 0.67 and p < 0.05 were used as screening criterion. CK0: no drought and no floods; CK1: drought without floods; CK2: no drought with floods; T1: abrupt drought-flood alteration.

### Gene functional description and Gene Ontology (GO) analysis

To annotate the function of T1 response proteins, proteins IDs were searched against the NCBI database (https://www.ncbi.nlm.nih.gov/) and/or the Uniprot database (http://www.uniprot.org/). Among the 522 DEPs between T1 and CK0, 244 up-regulated proteins and 231 down-regulated proteins could be annotated with functions, and nine up-regulated proteins and 38 down-regulated proteins remained uncharacterized. Among the 528 DEPs between T1 and CK1, 254 up-regulated proteins and 230 down-regulated proteins could be annotated with functions, and 13 up-regulated proteins and 31 down-regulated proteins remained uncharacterized. Among the 82 DEPs between T1 and CK2, 40 up-regulated proteins and 35 down-regulated proteins could be annotated with functions, and one up-regulated proteins and six down-regulated proteins remained uncharacterized. Summaries and expression patterns of proteins are listed in supplementary Excel sheets (sheet1 shows T1 vs CK0, sheet2 shows T1 vs CK1, and sheet3 shows T1 vs CK2).

To determine the cellular component, molecular function, and biological process categories of GO for T1 response proteins, we searched their protein IDs against the GO database (Mi and Al, 2013). 475 proteins could be annotated in certain GO categories between T1 and CK0. There were 14 significant GO terms for these differential proteins in the biological process, four significant GO terms for these differential proteins in the cellular component, and eight significant GO terms for these differential proteins in the molecular function, respectively (Fig. 7). The main biological functional categories were found to be metabolic pathways (20.03%), carbohydrate metabolism (16.03%), biosynthesis of secondary metabolites (14.19%), amino acid metabolism (8.51%), lipid metabolism (6.51%), biosynthesis of other secondary metabolites (5.51%), energy metabolism (4.84%), and carbon metabolism (4.01%). According to molecular functional properties, these proteins were mainly classified into folding, sorting, and degradation (3.17%), translation (2.17%), environmental adaptation (0.67%), signal transduction (0.50%), and transcription (0.50%) (see Fig. S1).

**Figure 7.**
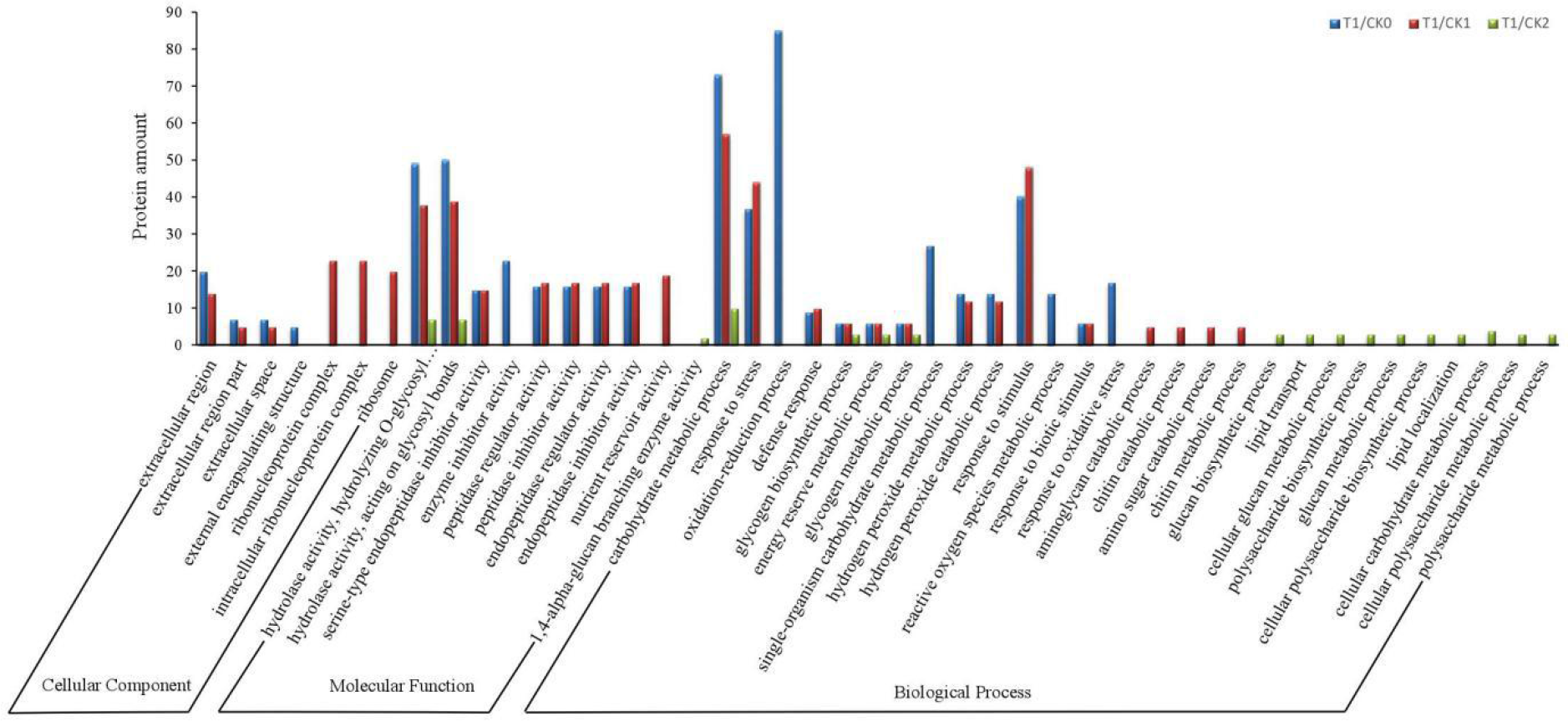
Gene Ontology (GO) enrichment analysis of differentially expressed proteins (DEPs) between T1 and CK0, CK1, and CK2. CK0: no drought and no floods; CK1: drought without floods; CK2: no drought with floods; T1: abrupt drought-flood alteration.

484 proteins could be annotated in certain GO categories between T1 and CK1. There were 14 significant GO terms for these differential proteins in the biological process, six significant GO terms for these differential proteins in the cellular component, and eight significant GO terms for these differential proteins in the molecular function, respectively (Fig. 7). The main biological functional categories were metabolic pathways (19.04%), biosynthesis of secondary metabolites (14.24%), carbohydrate metabolism (13.38%), amino acid metabolism (8.40%), biosynthesis of other secondary metabolites (5.15%), energy metabolism (5.15%), lipid metabolism (4.80%), and carbon metabolism (3.26%). According to the molecular functional properties, these proteins were mainly classified into folding, sorting, and degradation (4.97%), translation (3.77%), environmental adaptation (1.37%), signal transduction (0.51%), transcription (0.51%), and replication and repair (0.17%) (see Fig. S2).

75 proteins could be annotated in certain GO categories between T1 and CK2. There were 14 significant GO terms for these differential proteins in the biological process, zero significant GO terms for these differential proteins in the cellular component, and three significant GO terms for these differential proteins in the molecular function, respectively (Fig. 7). The main represented biological functional categories were metabolic pathways (18.92%), carbohydrate metabolism (16.22%), biosynthesis of secondary metabolites (13.51%), lipid metabolism (6.76%), metabolism of terpenoids and polyketides (5.41%), amino acid metabolism (4.05%), biosynthesis of other secondary metabolites (4.05%), energy metabolism (4.05%), carbon metabolism (4.05%), and metabolism of other amino acids (2.70%). According to molecular functional properties, these proteins could mainly be classified into folding, sorting, and degradation (10.81%), as well as transcription (2.70%) (see Fig. S3).

### KEGG pathways

A total of 522 DEPs were analyzed for the KEGG over-representation of pathways to obtain functional insights into the differences between the T1 and CK0 treatments. Of the 522 DEPs, 475 DEPs mapped onto the KEGG database. The significantly enriched KEGG pathways are listed in Table S2. The top three ranking canonical KEGG pathways are (in rank order): biosynthesis of secondary metabolites, metabolic pathways, and starch and sucrose metabolism, respectively. Detailed KEGG pathway information is listed in Table S2.

528 DEPs were analyzed for the KEGG over-representation of pathways to obtain functional insights into differences between T1 and CK1 treatment. Of the 528 DEPs, 484 DEPs mapped onto the KEGG database. The significantly enriched KEGG pathways are listed in Table S2. The top three ranking canonical KEGG pathways are (in rank order): biosynthesis of secondary metabolites, metabolic pathways, and diterpenoid biosynthesis, respectively. Detailed KEGG pathway information is listed in Table S2.

82 DEPs were analyzed for the KEGG over-representation of pathways to obtain functional insights into differences between T1 and CK1 treatments. Of the 82 DEPs, 75 DEPs mapped onto the KEGG database. The significantly enriched KEGG pathways are listed in Table S2. The top three ranking canonical KEGG pathways are (in rank order): starch and sucrose metabolism, alpha-Linolenic acid metabolism, and metabolic pathways, respectively. Detailed KEGG pathway information is listed in Table S2.

### Protein–protein interaction

All DEPs were uploaded into the SIMCA software package (version 14.0) to analyze protein interactions. This revealed that most enzymatic proteins and the biosynthesis of secondary metabolites, metabolic pathways, as well as starch and sucrose metabolism-related proteins interacted with T1 and CK0 treatments (see supplementary Fig. S4). Most enzymatic proteins and biosynthesis of secondary metabolites, and metabolic pathways-related proteins interacted with T1 and CK1 treatments (see supplementary Fig. S5). Most enzymatic proteins and starch and sucrose metabolism-related proteins interacted with T1 and CK2 treatments (see supplementary Fig. S6). Consistent with our metabolomics findings, GO analysis showed that the majority of proteins were involved in both carbohydrate metabolic process and hydrolase activity in the three comparison groups. Since the oxidation-reduction process was a significantly enriched GO between T1 and CK0, we exclusively focused on the carbohydrate metabolic process and oxidation-reduction process-related proteins at the proteomics level. Other proteins, such as signaling and molecular transport proteins, were excluded from further analysis.

## Discussion

The iTRAQ-labeled quantitative LC-MS/MS proteomics and LC-MS/MS metabolomics technology provides an effective method for the investigation of altered proteins and metabolites in plant cells during environmental stress (Frost *et al.*, 2015; Wu *et al.*, 2017). This study characterized the changes of the metabolome and the proteome of spike during the panicle differentiation stage in response to abrupt drought-flood alternation stress. This was achieved by comparing the relative expression patterns (treatment versus control) of the proteins and metabolites in rice. The aim was to obtain a more comprehensive understanding of the changes of the spike composition under abrupt drought-flood alternation stress and the molecular mechanism of rice yield decline. Further bioinformatics analysis showed the proteins that are important for the metabolic network and their regulation in response to abrupt drought-flood alternation. These findings provide the theoretical basis required for revealing the mechanism of abrupt drought-flood alternation yield formation.

### Response of metabolic regulation under drought-flood abrupt alternation stress

Photosynthesis is a biological process that is easily affected by drought and flood stress (Chaves *et al.*, 2009; Winkel *et al.*, 2016). Since water stress substantially decreases the CO2 assimilation via net reduction of the amount of ATP (Tezara *et al.*, 1999), it has been suggested that induction of ATP synthesis will assist in abiotic stress tolerance (Zhang *et al.*, 2008). In the present study, compared to CK0, CK1, and CK2, the net photosynthetic rate of T1 decreased at varying degrees (Fig. 1i). Furthermore, it is known that the KEGG pathway integration of *P*-values can be achieved based on differential proteomics and differential metabolomics, and carbon fixation in photosynthetic organisms pathway showed a significant difference of T1 treatment versus CK0 control (*P* < 0.01), T1 treatment versus CK1 (*P* < 0.01), and T1 treatment versus CK2 control (*P* < 0.05) (Table S2). The protein ribulose bisphosphate carboxylase (protein ID: A6N0B6), which is involved in carbon fixation in the photosynthetic pathway, significant down-regulated the T1 treatment versus CK0 control and T1 treatment versus CK1; no significant difference (*P* < 0.05) was found between T1 treatment versus CK2 control. This indicates that flood during the late stage can rapidly submerge plants, or fully submerged rice plants, resulting in a reduction of carbon dioxide assimilation; in response, the photosynthetic rate of rice under the condition of drought-flood abrupt alternation was inhibited. As a consequence, the amount of produced oxygen was reduced, and the respiration of rice was inhibited, thus limiting oxygen to the extent that oxidative phosphorylation no longer generates adequate ATP supplies and survival under low-oxygen conditions is directly linked to the ability to produce energy (ATP) under such circumstances. Low-oxygen tolerant plants (such as rice) are able to adequately respond to low oxygen by successfully remodeling their primary and mitochondrial metabolism to partially counteract the ensuing energy crisis (Shingakiwells *et al.*, 2015). The results show that the damage caused by abrupt drought-flood alternation on rice may mainly originate from late submergence damage.

Changes in carbohydrates are closely related to the plant response to drought and flood stress. Studies have shown that moderate drought stress is often caused by substances that increase the osmotic adjustment, such as carbohydrates, to enhance water retention and maintain cell expansion under stress conditions (Chaves *et al.*, 2009). Sugars act as osmoprotectants, help to maintain osmotic balance, stabilize macromolecules, abd provide an immediate energy source with which plants can restart growth (Yancey, 2005). Free proline, also an osmoprotectant, is accumulated in many plant species in response to environmental stress (Szabados and Savouré, 2010). The results of this study indicate that soluble protein content, MDA content, and free proline content under drought stress (CK1) during the panicle differentiation stage increased significantly (*P* < 0.01) compared to the CK0 control. Previous studies have shown that with further aggravation of water stress, the content of carbohydrates in plants can also be reduced (Pinheiro *et al.*, 2001). In this study, soluble protein content, free proline content, soluble sugar content under flood (CK2), and drought-flood abrupt alternation stress (T1) during the panicle differentiation stage decreased significantly (*P* < 0.01) compared to the CK0 control. To simulate the drought-flood abrupt alternation meteorological disasters in the Yangtze river valley in China, a total flooding time of 10 days was relatively long, and the degree of resulting stress was severe. As a result, the soluble sugar of rice leaves rapidly depleted to maintain the survival of rice, which may be the reason for the gradual depletion of energy reserves, such as starch and sucrose. At the same time, in T1, both the starch and sucrose metabolism pathways were significantly different compared to CK0, CK1, and CK2 (*P* < 0.01; see Table S2). The results of DEPs analysis indicate that the starch synthesis-related protein soluble starch synthase 1 (protein ID: A2Y9M4), the sucrose synthase associated protein sucrose synthase 1 (protein ID: A2XHR1), and the sucrose synthase 3 (protein ID: A2YNQ2) (see supplementary Excel sheets: sheet1 T1 vs CK0, sheet2 T1 vs CK1, and sheet3 T1 vs CK2) were significantly down-regulated, which supports the above viewpoint. DEPs of the T1 treatment versus CK0 control show that, osmotin-like protein (protein ID: A2WWU4) was significantly down-regulated. These results support each other with the significant decrease of soluble protein content in rice leaves under abrupt drought-flood alternation. Due to the reduction of energy reserve substances starch and sucrose, which failed to supply the glycolysis process with glucose, fructose, and other losses in a timely manner, and eventually the carbohydrate content gradually decreased and reduced carbohydrate osmotic adjustment function. The soluble protein content and free proline content were significantly decreased compared to the CK0 control (*P* < 0.01). Consequently, the energy supply during the following flood was not timely, the rice plants were in an energy starvation state, and the rice yield was reduced. KEGG pathway analysis showed that the carbon metabolism, glycolysis/gluconeogenesis, and pentose and glucuronate interconversions pathway are significant up-regulated in the T1 treatment versus the CK0 control (Table S2). The enhancement of glycolysis will result in acetyl CoA accumulation during the TCA cycle, and ultimately produce a large number of ATP in response to the abrupt drought-flood alternation stress; at the same time, mutual evidence supports the decline of soluble sugar content in physiological data. In addition, plant carbohydrates can be regulated as signaling molecules, interacting with other substances for signal networks (Candiano *et al.*, 2004; Rajjou *et al.*, 2006). It was found that cholic acid glucuronide was up-regulated in the T1 treatment group vs CK0 control (Table S1), thus confirming the occurrence of the glycolysis/gluconeogenesis pathway.

Lipid substances and fatty acids are another important energy storage substance in plants. Previously, Wang *et al.*, (2017) used C_13_ isotope calibration experiments to simultaneously calibrate glycerol (representing fatty acids) and glucose with the C_13_ isotope label in the experimental system. However, in the end, fatty acids rather than sugar were detected mainly in arbuscular mycorrhizal fungi, which for the first time rejected the accepted idea that sugar was the main carbon source from plants to mycorrhizal fungi. Furthermore, methods of genetics, molecular biology, and metabolic biology were used to study the findings. The fatty acid synthesis of plant hosts is necessary for the symbiosis of the arbuscular mycorrhizal fungi, and the fatty acids synthesized by plants can be directly transferred to mycorrhizal fungi. Previous studies in Arabidopsis reported that drought stress resulted in a decrease of the lipid content in the leaves and an enhancement of gene expression involved in the degradation of lipid substances (Gigon *et al.*, 2004). In this study, differential metabolites indicate that the contents of fatty acids, such as corchori fatty acid and conjugated linoleic acid (Table S1) were decreased under drought-flood abrupt alternation stress. According to the analysis of DEPs, it can be concluded that the abrupt drought-flood alternation stress inhibited the expression of stearoyl-[acyl-carrier-protein] 9-desaturase 2, Stearoyl-[acyl-carrier-protein] 9-desaturase 5 (see supplementary Excel sheets: sheet1 shows T1 vs CK0, sheet2 shows T1 vs CK1, and sheet3 shows T1 vs CK2), and the results of enhancing the expression of long chain acyl-CoA synthetase 4 and other enzymes involved in fatty acid degradation. Moreover, according to KEGG pathway enrichment analysis, fatty acid metabolism (*P* < 0.01), fatty acid degradation (*P* < 0.01), and fatty acid biosynthesis (*P* < 0.05) were significantly enriched (Table S2). The synthesis of fatty acids was enhanced to contribute to the promotion of the production and accelerate the rate of conversion between different substances. In addition, various lipid molecules produced via fatty acid degradation can act as important intracellular signaling molecules under various environmental stresses (Gigon *et al.*, 2004). Fatty acids and their derivatives play important signaling roles in plant defense responses (Jiang *et al.*, 2009) and transfer of these signal molecules contributes to a new abrupt drought-flood alternation adaptation reaction.

According to the analysis of metabolomics and proteomics data under the condition of abrupt drought-flood alternation, metabolites are mainly reflected in changes of sugar, fatty acids, and other molecules. Proteomics data mainly reflected the control of its metabolic process related enzymes and proteins changes. The data of these two dimensions showed that the energy metabolism was greatly affected under the condition of abrupt drought-flood alternation. The disturbance of the energy metabolism could not provide energy for growth and development of the rice panicle in time under the condition of abrupt drought-flood alternation, thus resulting in cell death and decreased rice yield.

### Damage mechanism of ROS under abrupt drought-flood alternation stress

In plants under flooding stress, the electron transport chains of mitochondria and chloroplasts is blocked, and the intracellular energy charge is reduced. All these factors could promote the production of reactive oxygen species (ROS) (Biemelt *et al.*, 1998). Then, the plant scavenging ROS system loses balance, the metabolism is blocked, and plants eventually die (Scandalios, 1993). Although the accumulation of ROS may be caused by oxidative stress, plants can adjust this through both their antioxidant enzyme system and their non-enzymatic system, such as non-enzyme scavengers and ROS scavenging enzymes, to resist the toxic effects of ROS under flooding stress. Consequently, plants alleviate the damage caused by ROS accumulation under flooding stress, which is beneficial for their survival after rice flooding stress (Ella *et al.*, 2003). POD, CAT, SOD, and glutathione-S-transferase usually act as ROS scavengers to reduce oxidative damage caused by oxidative stress in plants (Ahmed *et al.*, 2002; Wu *et al.*, 2013; Chen *et al.*, 2005). In this study, the T1 treatment versus CK0 control was investigated through differential proteomics significant enrichment biological process of GO in response to stress, oxidation-reduction process, defense response, hydrogen peroxide metabolic process, hydrogen peroxide catabolic process, response to stimulus, reactive oxygen species metabolic process, response to biotic stimulus, and response to oxidative stress process. Here, the scavenging of the ROS protein peroxidase 1 (protein ID: A2WPA9), peroxidase P7 (protein ID: B8B3L5), peroxygenase (protein ID: A2XVG1), cationic peroxidase SPC4 (protein ID: A2WZD6), peroxidase 15 (protein ID: A2Z4F1), L-ascorbate oxidase (protein ID: A2YE69), peroxidase 12 (protein ID: B8ARU4), cationic peroxidase SPC4 (protein ID: A2WZD9), peroxidase 51 (protein ID: A2XM89), cationic peroxidase SPC4 (protein ID: A2XZ79), probable glutathione S-transferase GSTU6 (protein ID: A2Z9K1), peroxidase 2 (protein ID: B8B5W7), and peroxidase 1 (protein ID: A2Y0P6) were significantly up-regulated (*P* < 0.05) (see supplementary Excel sheets: sheet1 shows T1 vs CK0, sheet2 shows T1 vs CK1, and sheet3 shows T1 vs CK2). Furthermore, the SOD, CAT, and POD activity of T1 in leaves increased significantly compared to CK0, the mutual evidence is supported by physiological data and proteomics data. Copalic acid and L-l-homoglutathione metabolites with antioxidant activity were significantly up-regulated in differential metabolomics (*P* < 0.005) and with the injury of the plant of phytomonic acid significantly down-regulated (*P* < 0.005). These results showed that rice exhibited more severe damage during drought-flood abrupt alternation, and ROS accumulated rapidly. To survive under stress, rice plants activate their antioxidant system and reduce the damage caused by abrupt drought-flood alternation. However, due to the long duration of drought-flood abrupt alternation (18 days), rice plants were more severely damaged. The anti-oxidant system of rice itself was not sufficient to remove the large amounts of ROS. Such excessive amounts of ROS attack the fatty acids in the cytoplasmic membrane, which then triggers the free radical chain reaction of fatty acids in the cytoplasmic membrane. Eventually, this leads to an increased permeability of the cytoplasm and electrolyte leakage, thus destroying the cell electrochemical balance, causing metabolic disorder, and cell death. In conclusion, this may be one of the important reasons for the decrease of rice yield due to abrupt drought-flood alternation.

## Supplementary data

**Figure S1.** Functional categories of the identified differentially expressed proteins (DEPs) between T1 (abrupt drought-flood alteration) and CK0 (no drought and no floods) treatment.

**Figure S2.** Functional categories of the identified differentially expressed proteins (DEPs) between T1 (abrupt drought-flood alteration) and CK1 (drought without floods) treatment.

**Figure S3.** Functional categories of the identified differentially expressed proteins (DEPs) between T1 (abrupt drought-flood alteration) and CK2 (no drought with floods) treatment.

**Figure S4.** Differentially expressed proteins (DEPs) protein–protein interaction analysis between T1 (abrupt drought-flood alteration) and CK0 (no drought and no floods) treatment.

**Figure S5.** Differentially expressed proteins (DEPs) protein–protein interaction analysis between T1 (abrupt drought-flood alteration) and CK1 (drought without floods) treatment.

**Figure S6.** Differentially expressed proteins (DEPs) protein–protein interaction analysis between T1 (abrupt drought-flood alteration) and CK2 (no drought with floods) treatment.

**Table S1.** Fold change (FC) for the abrupt drought-flood alternation responsive differential metabolites (DMs) between T1 and CK0, CK1, CK2.

**Table S2.** Enriched KEGG Pathways associated with differential metabolites (DMs) and differentially expressed proteins (DEPs).

**Excel Sheets S1 - S3.** Summaries of proteins in the different comparison groups

## Acknowledgements

This study was supported by the National Natural Science Foundation of China (Grant No. 31471441 and 30860136), the Jiangxi Science and Technology Support Project of China (Grant No. 2010BNA03600), and the Jiangxi Provincial Department of Education Project of China (Grant No. GJJ14283).

